# Tracking the emergence and dissemination of a *bla*_NDM-23_ Gene in a Multi-Drug Resistance Plasmid of *Klebsiella pneumoniae*

**DOI:** 10.1101/2022.07.05.498915

**Authors:** Neris García-González, Beatriz Beamud, Begoña Fuster, Salvador Giner, Ma Victoria Domínguez, Antonia Sánchez, Jordi Sevilla, Teresa Coque, Concepción Gimeno, Fernando González-Candelas, the Networked Laboratory for Antimicrobial Resistance Surveillance of Comunitat Valenciana (Spain)

**Author notes:** **Corresponding authors** Neris Garcia González, Fernando González-Candelas.

## Abstract

**Objectives:** Since the discovery of *bla*_NDM-1_, NDM beta-lactamases have become one of the most widespread carbapenemases worldwide. To date, 28 different NDM variants have been reported but some, such as *bla*_NDM-23_, have not been characterized in detail yet. Here, we describe the emergence of a novel *bla*_NDM-23_ allele from a *bla*_NDM-1_ ancestor and the multidrug resistant plasmid that has disseminated it through a *K. pneumoniae* ST407 clone in several Spanish hospitals.

**Methods:** Between 2016 and 2019, 1,972 isolates were collected in an epidemiological survey for ESBL-producing *Klebsiella pneumoniae* in the Comunitat Valenciana (Spain). Three carbapenem resistant strains failed to be detected by CPE screening tests. These isolates carried a *bla*_NDM-23_ gene. To characterize this gene, its emergence, and dissemination, we performed antimicrobial susceptibility tests, hybrid sequencing with Illumina and Nanopore technologies, and phylogenetic analyses.

**Results:** The MICs of the *bla*_NDM-23_ variant were identical to those of the *bla*_NDM-1_. The *bla*_NDM-23_ variant was found in 14 isolates in a 97 Kb non-mobilizable, multidrug-resistant plasmid carrying 19 resistance genes for 9 different antimicrobial families. In this plasmid, the *bla*_NDM-23_ gene is located in the variable region of a complex class-1 integron with a singular genetic environment. The short genetic distance between *bla*_NDM-23_-producing isolates reflects a 5-year-long clonal dispersion involving several hospitals and interregional spread.

**Conclusions:** We have characterized the genomic and epidemiological contexts in the emergence and community spread of a new *bla*_NDM-23_ allele in an MDR-plasmid of *Klebsiella pneumoniae*.

**Tweet:** Genomic, epidemiologic and phylogenetic analyses of the emergence of a new NDM allele provide information on the rapid changes underlying the spread of antimicrobial resistance genes and strains in *Klebsiella pneumoniae*.

**Importance:** At a time when antimicrobial resistance has become one of the biggest concerns worldwide, the emergence of novel alleles and extremely drug-resistant plasmids are a threat to public health worldwide. More so when they produce carbapenem resistance in one of the most problematic pathogens in clinical settings, such as *Klebsiella pneumoniae*. Here, we have used genomic epidemiology to describe the emergence of a novel NDM-23 allele and identify it in a MDR plasmid that has been disseminated through a *K. pneumoniae* ST407 clone in several hospitals in a Spanish region. By means of bioinformatic and phylogenetic analyses, we have been able to trace the evolutionary and epidemiological route of the new allele, the hosting plasmid, and the strain that carried both of them from Pakistan to Spain. A better understanding of the NDM-producing *K. pneumoniae* populations and its plasmids has made evident the spread of this clone through the region, enhancing the importance of genomic surveillance in the control of antimicrobial resistance.

## Introduction

The global dissemination of carbapenem-resistant *Enterobacteriaceae* has become a major threat to public health. Since the discovery of *bla*_NDM-1_ in a *Klebsiella pneumoniae* strain in 2008(1), NDM beta-lactamases have become one of the most widespread carbapenemases worldwide(2), due to their rapid evolution and dissemination via multidrug resistance (MDR) plasmids(3). *bla*_NDM_ genes are predominantly found in the *Enterobacteriaceae* family in which *K. pneumoniae* is the species with the highest frequency of these genes(4). Indeed, carbapenem-resistant *K. pneumoniae* represents the fastest growing antibiotic resistance threat in Europe, in terms of the number of infections and mortality(5). To date, 29 different NDM alleles have been reported, some of them showing an enhanced carbapenemase activity. Nevertheless, the dissemination, genetic context, and carbapenemase activity of some alleles remain unclear(6).

The first case of a *bla*_NDM_*-K. pneumoniae* producer in Spain was reported in 2012(7). Since then, and despite recent increases in prevalence, the total number of cases has remained low and they have mainly been associated with a few sporadic cases or small outbreaks. These outbreaks are primarily related to variants *bla*_NDM-1_ and *bla*_NDM-7_ carried by IncR, IncX3, IncN2, and IncFIB plasmids and associated with ST437, ST11, ST101, and ST147 strains(8, 9).

The Consorcio Hospital General Universitario de Valencia is a healthcare facility complex with a reference population of almost 360 000 inhabitants in Valencia, Spain. Since 2015, after an imported case of *bla*_NDM-1_-producing *K. pneumoniae*, the hospital noticed a substantial increase of this pathogen in the hospital(10). In 2018, an epidemiological surveillance program for *K. pneumoniae* was established in the Comunitat Valenciana (CV) region in Spain. Here, we report the emergence and dissemination of an ST437 clone carrying a *bla*_NDM-23_ in a MDR plasmid. In our work, we observed that this lineage was present in 5 hospitals in the region for 5 years. We also give a detailed characterization of the novel *bla*_NDM-23_ gene, its antimicrobial properties, and its immediate genetic context. A previous variant *bla*_NDM-23_ had been deposited as a complete CDS sequence in GenBank (accession MH450214.1). Still, no information about its genetic context, bacterial host, or antimicrobial activity has been published so far.

## Material and Methods

### Setting and sample collection, and bacterial isolation

During the period 2016-2019, we conducted the Networked Laboratory for Surveillance of Antimicrobial Resistance (NLSAR) of ESBL or carbapenemase-producing *K. pneumoniae* in the Comunitat Valenciana (Spain). Eight hospitals in the region participate in the Networked Laboratory. These are reference hospitals for more than 25 hospitals and healthcare centers. The first 30 clinical isolates of the month (if they had so) identified as ESBL or carbapenemase-producing *K. pneumoniae* in each hospital were included in the study. Additionally, 10 susceptible controls for each month and historical and retrospective samples from 2004 to 2017 were included. For epidemiological studies, relevant clinical data including isolate collection date, hospital ward, gender, and age were obtained from hospital records.

### Antimicrobial susceptibility and CPE screening test

Antimicrobial susceptibility tests were performed by the broth microdilution method using the MicroScan WalkAway (Beckman Coulter) automated system. The tested antibiotics varied depending on the hospital where each sample was collected. Susceptibility breakpoints were interpreted according to the recommendations of the EUCAST 2021 guidelines(11). Carbapenemase production was confirmed upon admission to the hospital using different methods.

### Transformation and conjugation assays

To evaluate the antimicrobial susceptibility of the relevant *bla*_NDM_ carrying isolates, we performed cloning and transformation assays. The complete sequences of *bla*_NDM_ genes were amplified as previously described(12). The PCR fragments were cloned using the TOPO TA Cloning kit (Life Technologies) and transformed into *E. coli* TOP10 background (Life Technologies). Transformants carrying the NDM genes were selected on plates containing kanamycin (50 μg/mL) and confirmed by PCR as above. Antimicrobial susceptibility testing of the transformants was performed using the MicroScan WalkAway (Beckman Coulter) automated system. For carbapenems, the minimum inhibitory concentration (MIC) was checked by the broth microdilution method. Three independent replicates for each isolate were obtained. Growth curves were represented using ggplot2(13).

To study the mobility of the plasmid carrying the *bla*_NDM_ gene, conjugation assays were performed using the clinical strains encoding carbapenemases as donors and a plasmid-free azide-resistant *E. coli* J53 strain as recipient. Three *K. pneumoniae* donors were studied: isolate 179KP-HG, carrying *bla*_NDM-1_, isolate 146KP-HG, carrying *bla*_NDM-23_, and isolate 262KP-HG, carrying *bla*_NDM-23_ and *bla*_OXA-48_. Donor and recipient cells were incubated separately overnight at 37°C with moderate shaking (150 rpm) in 1 mL of Luria-Bertani (LB) broth. Overnight cultures were ten-fold diluted and combined in a 1:1 donor-to-recipient ratio. Precipitated conjugation mixtures were cultured overnight at 37°C in a nitrocellulose membrane (0.22 µm pore diameter). Finally, recipients, donors, and transconjugants were selected on LB agar containing 0.5 μg/mL of ertapenem, 100 μg/mL of sodium azide, or both. Technical and biological replicates for each isolate were performed. Mobilization of the carbapenemase genes into *E. coli* J53 was confirmed by CHROMagar and PCR. PCRs for the *bla*_OXA-48_ gene were performed as previously described(14).

### Whole-genome sequencing and comparative analyses

Sequencing was performed in an Illumina NextSeq 500 platform using the Nextera XT library preparation kit. One isolate carrying *bla*_NDM-23_ (146KP-HG) and another carrying *bla*_NDM-1_ (179KP-HG) were also sequenced using Oxford Nanopore MinION technology. Sequencing reads and assemblies generated in this work are available at the ENA, project PRJEB37504 (ERS6077430 - ERS6077456).

To study the origin and evolution of *bla*_NDM-23_-carrying plasmids and their association with ST437 isolates, we analyzed the genetic context of all *bla*_NDM_ genes found in the NLSAR collection. Only isolates with a similar genetic context to that of the *bla*_NDM-23_ gene were analyzed in detail and kept for further results. To trace the evolutionary origin of these strains, we performed a comparative analysis including all ST437 genome sequences publicly available at GenBank (6/13/2020). Genomes with too many contigs (>500) were excluded. In addition, we included reads of ST437 isolates deposited at ENA from a previous study(9) about the emergence of NDM-producing *K. pneumoniae* and *Escherichia coli* in Spain. These reads were processed following the same workflow used for the reads obtained in our study.

Quality and filtering of short reads was assessed using FastQC(15) and Prinseq-lite(16). Reads with a mean quality below 25, a read length shorter than 60 bp, longer than 500 bp, or optical duplicates were removed. Also, the last three positions in the 3’ end and positions with a quality lower than 20 were trimmed.

Samples were *de novo* assembled using Unicycler(17) and gene annotation was performed with Prokka(18). QUAST(19) was used to assess the quality of all the assemblies. Detailed characterization was performed using Kleborate(20) and Kaptive(21). Additionally, ResFinder(22), PlasmidFinder(23), PLSDB(24), staramr(25), and BLAST were used to corroborate the antimicrobial resistance genes and plasmids detected. PHASTER(26) was used to analyze the presence of phages. To investigate the similarity between plasmids, BLAST(27), Prokka(18), and the R library genoplotR(28) were used. Plasmid relaxases, mobilization, and conjugation systems were searched using OriTFinder(29). Plasmid figures were drawn using the CGview server(30) and the genoplotR library.

Chromosome comparisons were made using Snippy(31). For this, the 146KP-HG hybrid assembled chromosome was used as reference. Minimum quality, minimum site depth for calling alleles, and minimum proportion for variant evidence were set to 60, 5, and 0.75, respectively. The quality of the genome sequence alignment was assessed using AMAS.py(32). We used snp-sites(33) to extract SNP positions from the whole genome alignment. Distance matrices were obtained using snp-dists (https://github.com/tseemann/snp-dists). Reference genome mapping coverage was calculated using the genomecov function of BEDTools(34). A maximum-likelihood phylogenetic tree was inferred from the genome alignment with IQTREE2(35) with the -TEST option to find the best fitting substitution model(36). Ultrafast bootstrap branch supports were assessed employing 1000 replicates(37). The ML tree was visualized using iTOL(38). To study the shared accessory genome, unmapped reads against the 146KP-HG chromosome were mapped against the plasmids found in this sample as above. The remaining unmapped reads were assembled using Unicycler and their gene content was analyzed using PROKKA and QUAST.

To avoid including highly divergent regions in the analyses, five complete prophages identified by PHASTER and an ICEkp yersiniabactin inserted in the reference chromosome were removed from the genome alignment. Temporal phylogenetic signal and estimation of the time to the most common recent ancestor were made using TempEST(39).

## Results

### Bacterial isolates, CPE screening tests, and antimicrobial susceptibility testing

We collected 1972 isolates under the NLSAR genomic surveillance program and detected 47 (2.31%) carrying NDM genes. Nevertheless, 3 of those 47 isolates yielded negative results in the CPE screening tests (Table S3). All of these isolates carried a *bla*_NDM_ gene with a novel variant, *bla*_NDM-23_. In total, 8 *bla*_NDM-23_-carrying strains were found, all of them belonging to ST437. These strains were non-susceptible to almost all the antimicrobials tested, except for colistin, tigecycline, and, occasionally, amikacin and fosfomycin (Table S2). Two ST437 isolates were sequenced by ONT, one carrying *bla*_NDM-23_ (146KP-HG) and *bla*_NDM-1_ (179KP-HG) the other.

### *bla*_NDM-23_ is located in a non-mobilizable, non-typable, multidrug resistance plasmid

The *bla*_NDM-23_ sequence differs from that of *bla*_NDM-1_ in one non-synonymous substitution at codon 101 (I101L). The MIC values for isolates carrying these variants were coincident for all the antibiotics, thus presenting the same antimicrobial susceptibility as *bla*_NDM-1_, which means low susceptibility to carbapenems and all beta-lactams (Table 1 and figure S1).

**Table 1.** Antimicrobial susceptibility testing for the clinical isolates and the TOP10 *E. fingers crossed coli* transformants carrying *bla*_NDM-1_ and *bla*_NDM-23_ genes.

|  | Clinical isolates |  |  |  | Transformants |  |  |  |
| --- | --- | --- | --- | --- | --- | --- | --- | --- |
|  | 179KP-HG (NDM-1) |  | 146KP-HG (NDM-23) |  | TOP10-NDM-1 |  | TOP10-NDM-23 |  |
| Ampicillin | >8 | R | >8 | R | >8 | R | >8 | R |
| Amoxicillin-clavulanate | >32 | R | >32 | R | >32 | R | >32 | R |
| Piperacillin-tazobactam | >8 | R | >16 | R | >16 | R | >16 | R |
| Cefuroxime | >16 | R | >8 | R | >8 | R | >8 | R |
| Cefotaxime | >8 | R | >32 | R | >32 | R | >32 | R |
| Cefepime | >16 | R | >8 | R | >8 | R | >8 | R |
| Aztreonam | >4 | R | >4 | R | ≤1 | S | ≤1 | S |
| Ertapenem<sup>a</sup> | >64 | R | >64 | R | >64 | R | >64 | R |
| Imipenem<sup>a</sup> | >64 | R | >64 | R | >64 | R | >64 | R |
| Meropenem<sup>a</sup> | >64 | R | >64 | R | >64 | R | >64 | R |
| Amikacin | ≤8 | S | ≤8 | S | ≤8 | S | ≤8 | S |
| Ciprofloxacin | >1 | R | >1 | R | ≤0.06 | S | ≤0.06 | S |
| Fosfomycin | ≤16 | S | ≤16 | S | ≤16 | S | ≤16 | S |
| Tigecycline | ≤8 | S | >2 | S | ≤1 | S | ≤1 | S |
| Trimethoprim | >4 |  | >4 |  | ≤2 |  | ≤2 |  |
| Cotrimoxazol | >4/76 | R | >4/76 | R | ≤2/38 | S* | ≤2/38 | S* |
| Colistin | ≤2 | S | ≤2 | S | ≤2 | S* | ≤2 | S* |

<sup>a</sup> The minimum inhibitory concentration (MIC) was checked by the broth microdilution method.

The hybrid assembly of isolate 146KP-HG revealed that the *bla*_NDM-23_ gene resides in a 97 kb plasmid hereafter referred to as plasmid p146KP-NDM23. In addition to p146KP-NDM23, this isolate carried three plasmids corresponding to a 118 Kb phage-like plasmid and 5 Kb and 3 Kb plasmids.

p146KP-NDM23 is a multidrug-resistant plasmid that, in addition to *bla*_NDM-23_, has 18 more genes associated with reduced susceptibility to nine different antimicrobial families, including beta-lactams, sulfonamides, aminoglycosides, trimethoprim, fluoroquinolones, tetracycline, chloramphenicol, fosfomycin, tunicamycin, and rifampicin. All these genes are inserted in a large multiresistant region (MRR) that contains different putative transposable modules (Figure 1A), which have been previously detected in IncFII plasmids(40, 41). They include the gene array comprising *bla*_OXA-1_, a truncated *catB* gene, and *aac(6’)-Ib-cr* flanked by two copies of IS*26*; the tetracycline resistance gene, *tetA*, associated with the Tn*As1* transposase; genes *aac(3)-IIa* and *tmrB* linked to a copy of IS*26*; and a ΔTn*3* (*bla*_TEM-1B_) with the transposase (*tnpA*) disrupted by an IS*Ecp1*-*bla*_CTX-M-15_, with *sul2, strA*, and *strB* genes located downstream a *bla*_TEM-1B_.

**Figure 1.**
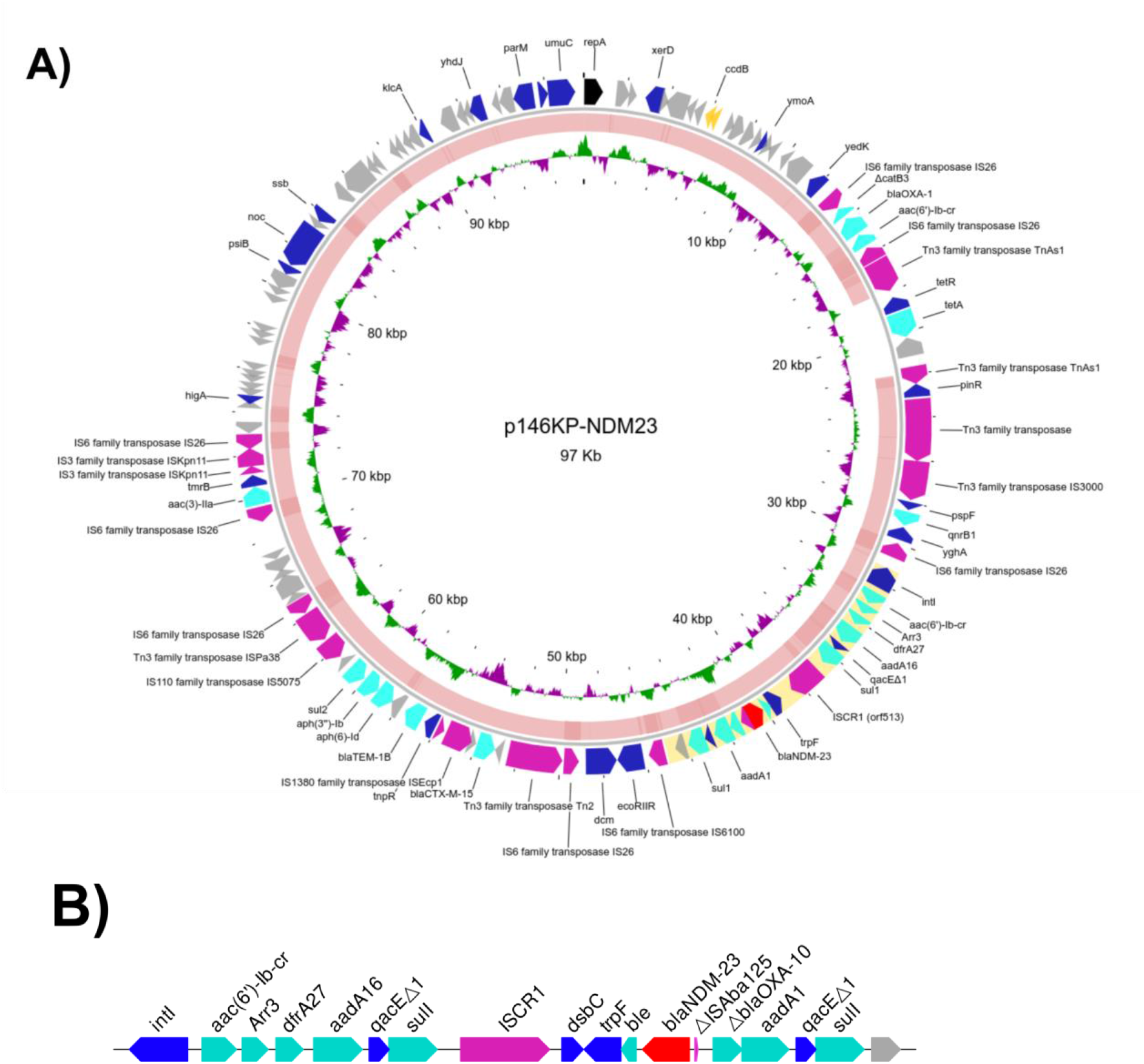
A) Gene content and structure of plasmid p146KP-NDM23. The *bla*_NDM-23_ embedded in the second variable part of a ISCR1 complex class 1 integron is highlighted in yellow. The middle ring shows the BLAST comparison between p146KP-NDM23 and p179KP-HG. The inner ring shows the GC skew of the plasmid. B) ISCR1 Complex class 1 integron were the *bla*_NDM-23_ gene is embedded.

The *bla*_NDM-23_ gene is in a complex class 1 integron associated with the ISCR1 (Figure 1B), as previously described(4). In the first variable region of the integron, we found *acc(6’)-Ib-cr, arr3, drfA27*, and *aadA16* gene cassettes. Then, the ISCR1 region comprised an *aadA1* gene, a truncated *bla*_OXA-10_ gene, and a truncated IS*Aba125* downstream the *bla*_NDM-23_ gene while upstream we found a *ble*_MBL_ gene, a phosphoribosyl anthranilate isomerase gene (*trpF)*, and an oxidoreductase *dsbC*.

The type of plasmid p146KP-NDM23 could not be identified either by replicon (Inc) or relaxase (MOB) typing. Although a BLAST search with RepA returned 24 plasmids with 100% identity to the query sequence (Table S4), no incompatibility type could be assigned to any of them. Apart from the *repA* gene, no homology was found for the rest of the sequence of p146KP-NDM23 and any of the 24 plasmids. We could not detect any relaxases or transfer genes in p146KP-NDM23. Thus, mobilization of p146KP-NDM23 could only occur in *trans* with help of co-resident plasmids; nevertheless, conjugation assays showed no mobilization to the receptor cells.

### Origin of the p143KP-NDM23 plasmid

When comparing the p146KP-NDM23 sequence with the aforementioned databases, the most similar plasmid to p146KP-NDM23 in terms of identity and shared genes (100% coverage and identity, table S5) was pKDO1 (accession JX424423). Plasmid pKDO1 was isolated from an ST416 *K. pneumoniae* strain in the Czech Republic(42). pKDO1 contains all the genes found in plasmid p146KP-NDM23 with rearrangements, but not the ISCR1 complex class 1 integron where the *bla*_NDM-23_ gene is embedded (Figure 2).

**Figure 2.**
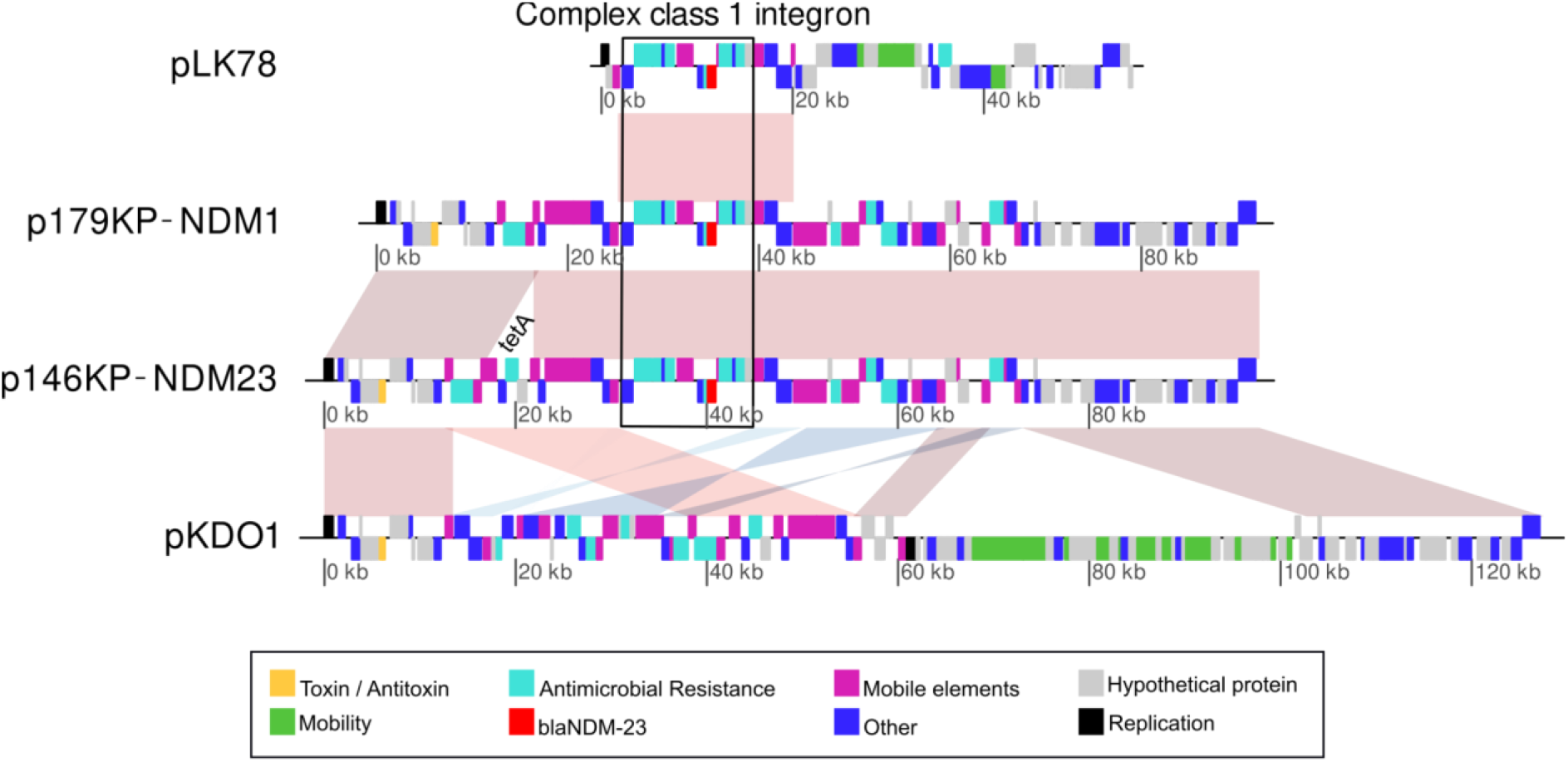
Structure comparison of plasmids pLK78, p146KP-NDM23, pKP179-NDM1, and pKDO1. pLK78 shares similarity with the integron sequence found in pKP146-NDM23, pKP179-NDM1, while pKDO1 shares similarity with the backbone of those plasmids.

However, complex class 1 integrons and ISCR1 associated with *bla*_NDM-23_ genes have been described previously (4). The structure around the *bla*_NDM-23_ gene in p146KP-NDM23 is not the usually conserved structure found around *bla*_NDM_ genes. In this case, we found the presence of a truncated *bla*_OXA-1_ downstream the *bla*_NDM_-ΔIsaba125. This genetic environment has been described previously in different plasmids (Table S6) but carrying the *bla*_NDM-1_ variant. An IncN3 plasmid, denoted as pLK78 (accession KJ440075), was found to carry the most similar environment to that of *bla*_NDM-23_ in p146KP-NDM23. Plasmid pLK78 was isolated in 2012 in Taiwan and collected from a *K. pneumoniae* strain(43). Sequence comparison between plasmids pLK78 and p146KP-NDM23 showed that they only have the ISCR1 complex class 1 integron with the *bla*_NDM_-coding in common, with a 99.9% of identity and 100% of coverage, while the rest of the plasmid has no similarity. The identity between these two integrons is not perfect because of the single substitution that differentiates the *bla*_NDM-1_ variant from *bla*_NDM-23_.

We detected the presence of the p146KP-NDM23 plasmid, the ISCR1 complex class 1 integron, and the previous pKDO1 and pLK78 plasmid sequences with coverages of 95.1%, 100%, 99.83%, and 92.39%, respectively, in the genome of a *K. pneumoniae* isolate named PN4(44). PN4 is a strain involved in a multispecies outbreak in 2010 in Pakistan. It was not possible to decipher the structure of each mobile genetic element in this strain because PN4 is available only as a draft genome assembly. Nonetheless, we have found a possible origin where the plasmid could have formed as we have identified all the necessary structures in a single isolate.

When we compared plasmid p146KP-NDM23 with the other strain sequence obtained from long reads in this work, 179KP-HG, we found a plasmid almost identical to p146KP-NDM23 but carrying the *bla*_NDM-1_ variant and lacking a TnAs1 transposable element carrying a *tetA* resistance gene (Figure 1A). We named this plasmid p179KP-NDM1.

### The recent evolutionary history of *bla*_NDM-23_-producing isolates is associated with ST437

All *bla*_NDM-23_ genes detected in the surveillance program were found in ST437 strains (Table S1). Thus, to analyze the epidemiological and evolutionary dynamics of the clonal background of the new *bla*_NDM-23_ gene and its plasmid, we included all the ST437 isolates collected in the NLSAR surveillance program that carried NDM genes (8 isolates) or not (14 isolates). We also included all the ST437 genomes from the RefSeq (121 isolates, accessed on 27/04/2021) (Table S7).

Reads from all the samples were mapped to the 146KP-HG closed chromosome derived from the hybrid assembly. The mapping coverage values ranged from 91.6% to 99.9% with an average of 96.27% (Table S8). Integrative mobile genetic elements were identified and removed from the final alignment: five complete prophages identified by PHASTER (Table S9) and an ICEkp yersiniabactin. This resulted in a sequence alignment of 143 complete genomes spanning 5,063,248 bp of which 5,960 corresponded to variant positions (SNPs).

The phylogenetic analysis revealed that the ST437 global population collected in this work is divided into three main groups corresponding to different capsular types. Only the isolates belonging to capsular type KL36 (Figure 3) were kept for further analysis as all the *bla*_NDM-23_-producers were found in KL36 isolates. The distance matrix for all the isolates is shown in Table S10 and the ML tree can be found in Figure S2. The model used to reconstruct the phylogenetic tree was GTR+F+I+G4.

**Figure 3.**
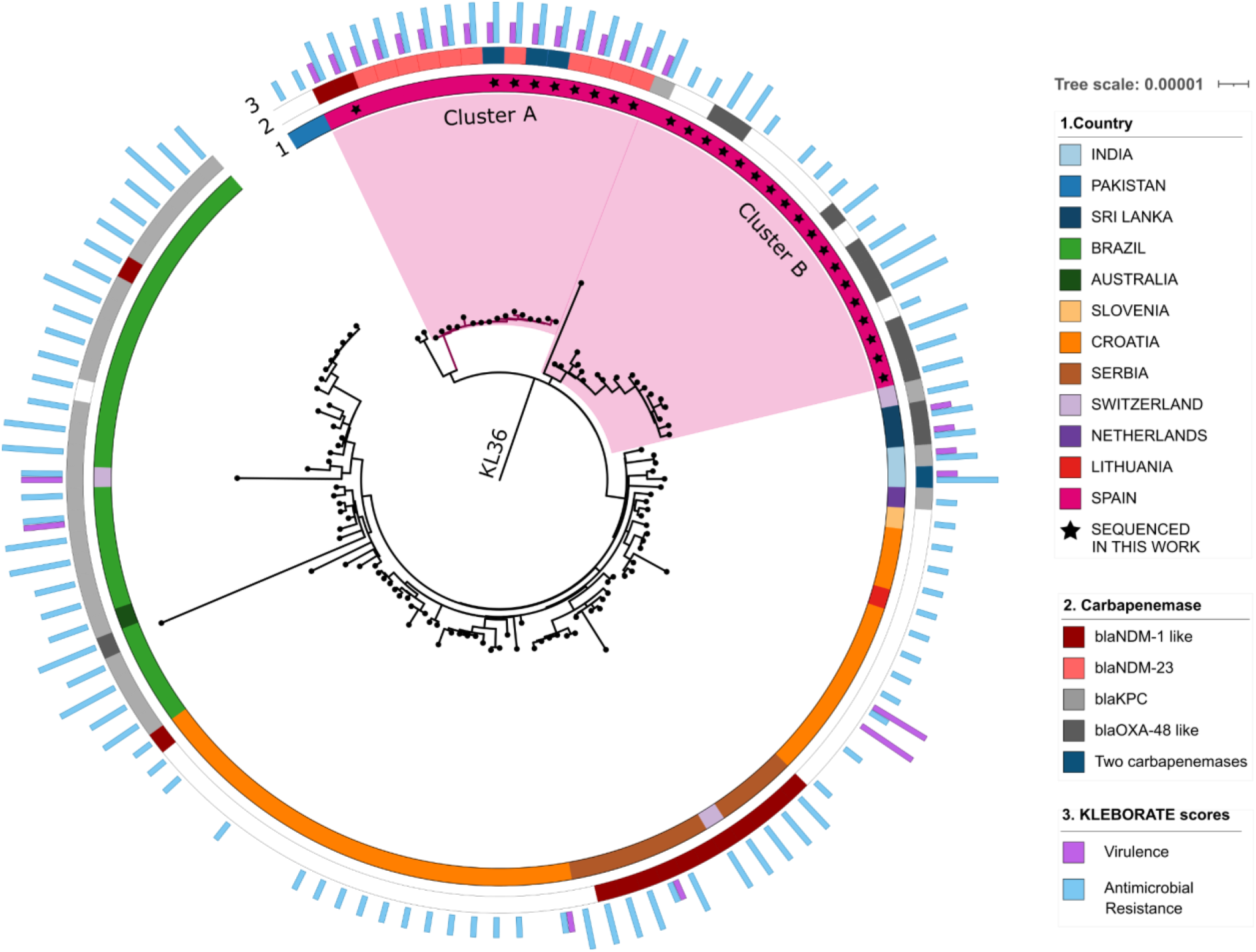
Whole genome maximum likelihood tree of ST437 – KL36 isolates. Spanish isolates are highlighted in light pink. Colors in the inner ring (1) indicate the country of origin of each sample. A star inside this ring indicates that the isolate has been sequenced in this work. Colors in the center ring (2) represent carbapenem resistant genes. In the outer ring (3), blue bars show the resistance score while purple bars show the virulence score obtained from Kleborate (20). The scale represents nucleotide divergence as substitutions per site.

Within KL36, all Spanish isolates were clustered in cluster A or cluster B. *bla*_NDM-23_-carriers were found only in cluster A (Figure 3). This cluster contains 16 isolates, all of them with a *bla*_NDM_ gene in the accessory genome and collected in Spain. Nine isolates were collected in the NLSAR surveillance program while seven isolates were downloaded from the database. These database genomes were isolated in 2016 (9, 45) and all of them carried the *bla*_NDM-23_ variant. This implies that the *bla*_NDM-23_ variant was already in Spain before we noticed it in the surveillance of NLSAR. The low number of SNPs among cluster A genomes (from 0 to 30, with an average of 11) and the highly conserved accessory genome of this clade suggest a recent common ancestry. Temporal analysis showed a very recent time to the MCRA of cluster A in mid-year 2013 (R^2^ = 0.47).

The remaining isolates collected in Spain and the NLSAR surveillance program grouped in cluster B. Although cluster B contains NLSAR isolates collected from the same hospitals and dates as those in cluster A, the phylogenetic tree shows that these clusters likely represent two different introductions of ST437-KL36 in the region and that the tMRCA of cluster B isolates occurred around 1999 (R^2^ = 0.60). This is also supported by the on average 141 SNPs (range 84-275) and the very different accessory genomes between cluster A and cluster B (range 224 - 1281 genes). The phylogenetic temporal analysis with TempEst showed that the ancestor of clusters A and B occurred in 1996 (R^2^=0.35). Hence, the Spanish non-NDM-producing ST437 isolates sequenced in this work were not the founder population of the NDM23-producing isolates.

The phylogeny (Figure 3) shows that the closest isolates to cluster A are two samples from Pakistan collected in 2011 (GCA_900181335 and GCA_900181325). These isolates show an average difference of 146 SNPs (range 131-157) with cluster A isolates. These SNPs are dispersed over the whole genome and they do not cluster in one or a few loci. Hence, we can eliminate the possibility that they have arisen from recombination or horizontal gene transfer. In addition, the Pakistan and cluster A samples share an *E. coli* phage, phiV10 (accession NC_007804), inserted in the chromosome that is not present in any of the other samples, further supporting their common ancestry. TempEst analysis estimated the tMCRA between the Pakistan and cluster A samples in 1999 (R^2^=0.78).

### Transmission of plasmid p146KP-NDM23 carrying *bla*_NDM-23_ by recent clonal dissemination

Clade A comprises all *bla*_NDM-23_ isolates, and all of them carry this gene in p146KP-NDM23. All the antimicrobial resistance determinants found in these strains were present in the p146KP-NDM23 plasmid. No other antimicrobial resistance genes or relevant mutations (OmpK, parC,…) were found in the chromosome or other plasmids of these strains except for intrinsic resistance genes (*bla*_SHV-11_, *FosA, OqxA*, and *OqxB*). In addition, this clade shares a yersiniabactin virulence factor (*ybt9*) in its chromosome. Additionally, we found that both *bla*_NDM-23_ and *bla*_OXA-48_ genes were present in 3 isolates.

Supported by the minimal number of differences found between the chromosomes of clade A isolates, the identical accessory genome, and the date of the tMCRA of the clade (mid-2013), we assumed that clade A represents a case of recent clonal dissemination. In addition to hospital H0 in Madrid (GCA_011684095) and hospital H1 in Valencia, this dissemination also affected at least three different hospitals in the Comunidad Valenciana, spanning from 2016 until at least 2019, when our sampling concluded. This clade comprises two basal isolates (GCA_011684095 and 179KP-HG) carrying the carbapenemase gene *bla*_NDM-1_ and 14 isolates that carry the novel *bla*_NDM-23_ allele. The index case of the dissemination of this clone, GCA_011684095, is the most basal genome of cluster A and was also the index case of a multi-clonal and multispecies outbreak detected in hospital H0 in Madrid(45). This patient was a 36-year-old man, native of Pakistan with residence in Valencia. The patient had received healthcare assistance in Pakistan following a traffic accident in November 2014. On 26 August 2015, upon return from Pakistan, he was admitted to the neurosurgery ward in H0 in Madrid. In December 2015, the patient was transferred to Valencia and admitted to hospital H1. A few months later, this patient went through the emergency ward in H2 and was admitted to the general ward. This explains the fast spread of the lineage through the region but also points to Pakistan as the possible origin of the strain causing this dissemination event.

The *bla*_NDM-23_ mutation had to occur between the arrival of the clones at the hospital and the first detections of NDM-23 genes in the hospital. The first clone of clade A carried *bla*_NDM-1_. After that, the p146KP-NDM23 plasmid remained genetically stable over time. We found 100% identity for p146KP-NDM23 plasmid among all the isolates but with an average coverage of 97.5%, ranging from 84% to 100% (Table S11, Figure 4). The plasmid has accrued several structural changes, mainly deletions (Figure 4). The accessory genome of the corresponding samples did change during the dissemination process, with gains and losses of complete mobile genetic elements. For instance, samples 262KP-HG, HAV1_09, and KP-HGUA01_12, have an additional IncL plasmid harboring a *bla*_OXA-48_ gene. Contrarily, some samples have lost a few complete plasmids: for instance, sample KP-HGUA01_12 lacks 118 kb, 5 kb, and 3 kb plasmids, and samples HLF2_50, HGV2_363 and HGV2_364 lack plasmid 118Kb.

**Figure 4.**
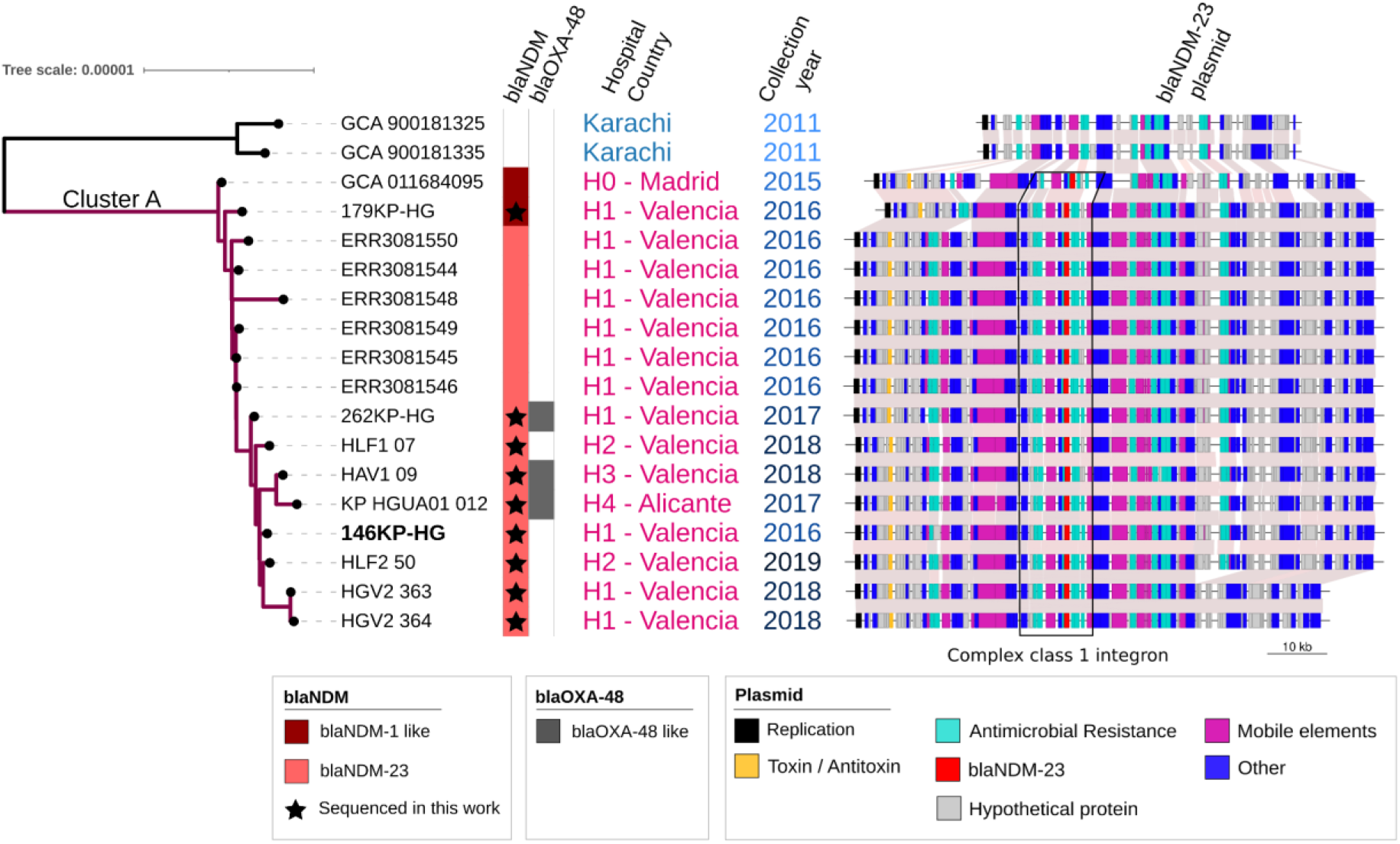
Dissemination of the ST437 clone carrying the *bla*_NDM23_ gene involves 5 different hospitals along 5 years. Isolates sequenced in this work are marked with a star inside the *bla*_NDM_ strip. For each isolate, the structural variants of plasmid p146KP-NDM23 is shown.

## Discussion

The dissemination of *bla*_NDM_ genes in a broad range of Gram-negative bacteria, such as *K. pneumoniae*, via MDR plasmids has established NDM as a global major public health threat(3). The evolutionary and molecular mechanisms acting on plasmids and the genes they contain are well understood at a general scale, but there is little detailed information on how they operate at short time scales.(46) A better understanding of these processes is thus essential for controlling and limiting the spread of ARGs.

Here, we have described the emergence and dissemination of a multi-drug-resistant plasmid associated with clonal dissemination of a *K. pneumoniae* ST437 strain carrying a new *bla*_NDM-23_ carbapenemase gene and 18 more antimicrobial resistance genes. We have also detected when and where a point mutation in the plasmid sequence produced a change from a *bla*_NDM-1_ allele to *bla*_NDM-23_ during the dissemination of this clone.

The multi-drug-resistant plasmids in which the *bla*_NDM-1_ and *bla*_NDM-23_ genes were identified were named pKP179-NDM1 and pKP146-NDM23, respectively. These plasmids contain the *bla*_NDM_ variants in a complex class 1 integron associated to ISCR1, in addition to 18 genes that confer resistance to nine different antimicrobial families. These plasmids were found to be very similar to plasmid pKDO1, described in the Czech Republic in 2009(42) while the complex class 1 integron was identified in plasmid pLK78, isolated in Taiwan in 2012(43). Contigs from the *K. pneumoniae* strain PN4 genome, isolated in Pakistan in 2012, included all the plasmids (pKP146-NDM23, pLK78, and pKDO1). Hence, we hypothesize that the pKP146-NDM23 originated in this lineage. We found a conjugation module in pKDO1 but not in pKP179-NDM1 nor pKP146-NDM23, suggesting that the latter plasmids lost their mobilization capabilities before their mobilization to the ancestor of Clade A.

Although conjugation assays for p146KP-NDM23 were negative, our bioinformatic analyses show the presence of the complex class 1 integron in different plasmids (Table S6). This might be explained either because the integron is flanked by IS26 transposases, which could mediate its transposition, or, although it has not been proven experimentally, by the independent mobilization of ISCR1 elements(47). Nevertheless, further work will be necessary to understand the mobilization of these genetic elements in different strains and species.

The close similarity of the chromosome sequences of isolates in clade A (Figure 4) and two isolates from Pakistan, the origin of the p146KP-NDM23, and the epidemiological link between the index case in Spain and that country strongly suggest the origin of cluster A sequences in Pakistan and not in other ST437 isolates previously found in Spain (cluster B). Moreover, the results indicated a recent emergence and clonal expansion of clade A in the Comunidad Valenciana.

Whereas the outbreak produced in 2015 at H0 in Madrid was contained at a single hospital, the spread of this clone throughout the Comunidad Valenciana in the following years, until at least 2019, represents the first case of interhospital and community spread of this clone in our country. In some of these hospitals, including H0(45), the *bla*_NDM-23_ allele was not detectable by some carbapenemase tests, and its actual spread might be underestimated. Remarkably, along its recent and short (5-year) dissemination, the plasmid p146KP-NDM23 has shown complete conservation at the nucleotide level but several differences at the structural level.

The *K. pneumoniae* clone that is disseminating this plasmid carries a yersiniabactin virulence factor, which encodes for an iron-scavenging molecule that enhances its capacity to cause disease associated with bacteremia and tissue-invasive infections(48). This combination, the limited number of effective therapeutic options available to treat infections caused by this clone, and the increasing number of infections in this region make this clone and its plasmid of particular concern for local and regional public health.

An important goal of our surveillance program is to inform about new threats and facilitate improved intervention strategies for controlling their spread. Infectious disease specialists should be aware of the possibility of finding this clone in their hospitals. Remarkably, carbapenemase detection was negative when some of these patients were tested for carbapenemase colonization(45). This highlights the importance of updating diagnostic techniques in hospitals to prevent further dissemination of this clone and detect this carbapenemase. As a non-mobilizable plasmid, it can only be transmitted by clonal dissemination, and hospital control measures will have to focus on timely detection of *bla*_NDM-23_ and finding and removing the source of this clone from hospitals.

## Acknowledgements

We are thankful to Dr. Waylan for kindly providing the genome assembly of PN4.

## Funding

This research was supported by project BFU2017-89594R (MICIN, Spanish Government) and by the European Union through the Operational Program of European Regional Development Fund (ERDF) of Valencia Region (Spain) 2014-2020. Additional support provided by the Network Research Centre for Epidemiology and Public Health (CIBERESP).

## List of members of the Networked Laboratory for Surveillance of Antimicrobial Resistance of the Valencian Community

Neris García-González and Fernando González-Candelas. Joint Research Unit Infection and Public Health FISABIO-University of Valencia, Institute for Integrative Systems Biology (I2SysBio). CIBER in Epidemiology and Public Health. Valencia. Spain.

Begoña Fuster, Nuria Tormo, Carme Salvador, and Concepción Gimeno. Microbiology Service. Consorcio Hospital General Universitario. Valencia, Spain.

Victoria Domínguez. Microbiology Service. Hospital Arnau de Vilanova. Valencia, Spain.

Salvador Giner. Microbiology Service. Hospital Universitario y Politécnico La Fe. Valencia, Spain.

Javier Colomina and David Navarro. Microbiology Service. Hospital Clínico Universitario. Valencia, Spain.

Llúcia Martínez. Sequencing Platform. FISABIO. Valencia, Spain.

Antonio Burgos and Olalla Martínez. Microbiology Section. Hospital La Ribera, Alzira, Spain.

Barbara Gomila-Sard and Rosario Moreno. Microbiology Service. Hospital General Universitario. Castelló, Spain.

Inmaculada Vidal, Antonia Sánchez and Juan Carlos Rodriguez. Microbiology Service. Hospital General Universitario. Alacant, Spain.

Victoria Sánchez-Hellín. Microbiology Service. Hospital General Universitario. Elx, Spain.

## Transparency declarations

None of the authors of this manuscript declares any conflict of interest.

## Figure Captions

**Supplementary Figure 1**. Comparative dynamics of growth under different concentrations of carbapenem antibiotics for 3 clinical isolates and 2 transformed E. coli TOP10 strains with NDM1 and NDM3.

**Supplementary Figure 2**. Maximum Likelihood phylogenetic tree from complete genome sequences of ST437 isolates included in this study.

